# Exploring intra-species diversity through non-redundant pangenome assemblies

**DOI:** 10.1101/2022.03.25.485477

**Authors:** Fernando Puente-Sánchez, Matthias Hoetzinger, Moritz Buck, Stefan Bertilsson

## Abstract

At the genome level, microorganisms are highly adaptable both in terms of allele and gene composition. Such heritable traits emerge in response to different environmental niches and can have a profound influence on microbial community dynamics. As a consequence of this, any individual genome or clonal population will contain merely a fraction of the total genetic diversity of any operationally defined “species”, with the collective from that group presenting the broader genomic diversity known as the pangenome. Pangenomes are valuable concepts for studying evolution and adaptation in microorganisms, as they partition genomes into core regions (present in all the genomes, and responsible for housekeeping and species-level niche adaptation) and accessory regions (present only in some genomes, and responsible for ecotype divergence).

Here we present *SuperPang*, an algorithm capable of producing pangenome assemblies from a set of input genomes of varying quality, including metagenome-assembled genomes or MAGs. *SuperPang* runs in linear time and its results are complete, non-redundant, preserve gene ordering and contain both coding and non-coding regions. Our approach provides a modular view of the pangenome, identifying operons and genomic islands, and allowing to track their prevalence in different populations. We illustrate our approach by analyzing the intra-species diversity of *Polynucleobacter*, a clade of ubiquitous freshwater microorganisms characterized by their streamlined genomes and their ecological versatility. We show how *SuperPang* facilitates the simultaneous analysis of allelic and gene content variation under different environmental pressures, allowing us to study the drivers of microbial diversification at unprecedented resolution.

## INTRODUCTION

Over the last decades, advances in sequencing and bioinformatics methods have led to a continuous increase in the resolution at which complex microbiomes can be analyzed. In line with this, the focus in many areas of microbiology research has gradually shifted from studies of individual populations or species to encompass entire microbial communities by means of large scale sequencing approaches (Inkpen *et al*., 2017). This has increased our understanding of how metabolic functions are distributed within communities and the ecological interactions that underpin their dynamics and change over time, space, and along environmental gradients (Louca *et al*., 2018; Galand *et al*., 2018). However, these advances also showed that variability was prevalent at all taxonomic scales, including striking differences in functional properties and environmental partitioning between microorganisms belonging to the same species (Larkin &Martiny, 2017).

Microbial species are comprised of multiple strains that share a core genome and differ in the accessory genome. Recently the term “pangenome” has been coined, referring to the combined pool of genes found in the known strains of a species and this pangenome can be partitioned into a core and accessory component (Tettelin *et al*., 2005). Variations in the accessory genome are assumed to be responsible for niche differentiation and allow the discrimination of ecologically meaningful subpopulations (Cohan, 2001; Coleman &Chrisholm, 2007; Koeppel *et al*., 2013). This variation, often referred to as “intra-species diversity” or “microdiversity”, have profound consequences for the functioning of microbial communities (Fuhrman &Campbell, 1998; García-García *et al*., 2019). However, its extent and specific roles remain poorly characterized due to the difficulty in obtaining high quality genomes from relevant environmental populations. The description of microbial species has traditionally been dependant on culturability, which greatly limited our view of microbial biodiversity (Sanford *et al*., 2021). However, recent methodological advances in sequencing capacity and bioinformatic tools have enabled reconstruction of near-complete microbial genomes from shotgun metagenomic data (Metagenome-Assembled Genomes, or MAGs), and also group them based on their average nucleotide identity (ANI) into metagenomic operational taxonomic OTUs (mOTUs) which correspond to species-level clusters from a purely genomic viewpoint (Sunagawa *et al*., 2015; Richter and Rosselló-Móra, 2009; Jain *et al*., 2018).

This wealth of culture-independent genomic data has rendered the concept of a single reference genome obsolete (Coleman &Korem, 2021), spurring the development of tools aiming to capture the full complexity of microbial pangenomes such as Roary (Page *et* al., 2015), PPanGGOLiN (Gautreau *et al*., 2020), mOTUpan (Buck *et al*., 2021) or PanACoTA (Perrin &Rocha, 2021). These tools work by generating clusters of orthologous genes from the input genomes, and then rely on different statistical approaches to classify each cluster these as part of the core or accessory genome. A disadvantage is that this approach relies on external prediction software such as Prodigal (Hyatt *et al*., 2010) for identifying genes in the input genomes. This is a straightforward task for genes encoding proteins or ribosomal/transfence RNAs, but becomes increasingly difficult for non-coding regions which might nonetheless mediate essential functions (Rogozin *et al*., 2002). Another shortcoming is that information on synteny is in principle lost when breaking the genome into separate features, although methods such as Roary and PpanGGOLin do keep track of gene neighborhood information when classifying clusters into the core or accessory gene sets.

An alternative to this is to directly build assembly graphs from sequence data in order to represent the pangenome (Brown, et al., 2020; Quince *et al*., 2021). Notably, the STRONG pipeline (Quince *et al*., 2021) aims to build a coassembly graph from different samples using metaSPAdes (Nurk *et al*., 2017), after which it performs binning and haplotype resolution. This method is particularly well suited for longitudinal or time-series metagenomic studies, but does not allow includsion of genomes obtained from other sources, such as isolates or Single-Cell Amplified Genomes (SAGs), or even MAGs obtained from other studies. Furthermore, the high memory usage of metaSPAdes makes it impractical for assembling very large datasets (Van der Walt *et al*., 2017), constraining the number of samples that can be analyzed with a coassembly-based approach.

## SOFTWARE DESCRIPTION

Here we present the *SuperPang* software to generate non-redundant pangenome assemblies from multiple input genomes from the same species/mOTU. The input genomes can come from isolates, SAGs or MAGs indistinctively, and genomes from different sources and different qualities can be combined in the same analysis. **Figure 1** summarizes the functioning and results of *SuperPang*, and how they compare to those obtained with raw pangenomes or single reference genomes.

**Figure 1.**
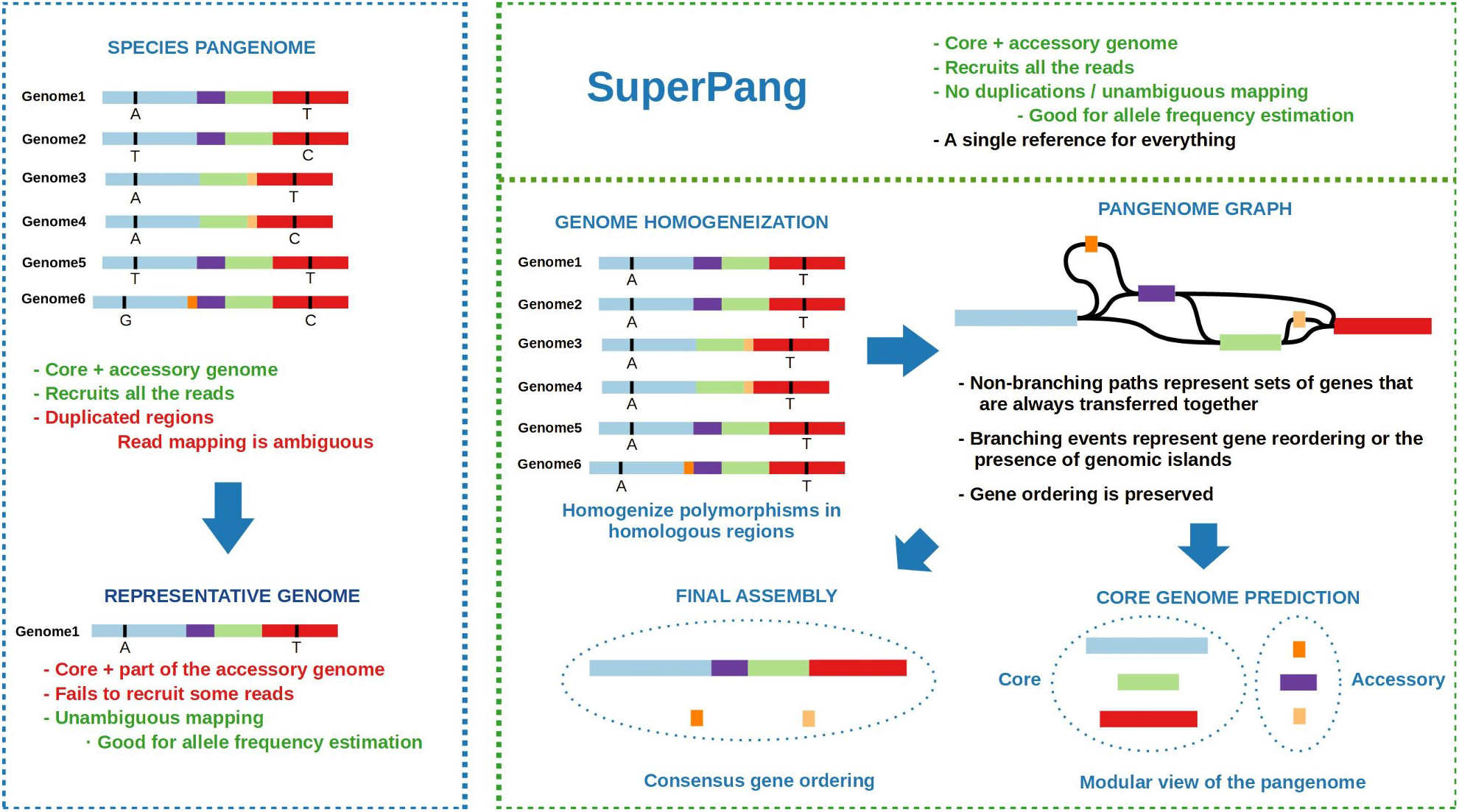
Strategies for analyzing the genetic diversity of microbial species. **Upper left:** the use of all the available genomes from the species of interest allows to analyze the totality of the accessory genome, but many regions will be redundant, resulting in ambiguous read mapping. **Lower left:** choosing a single reference genome for mapping will result in non-ambiguous mappings, but reads originating from the missing accessory genes will not be recruited. **Right:** *SuperPang* will produce a complete and non-redundant pangenome assembly partitioned into core and accessory components that preserves gene ordering and can be used for different applications.

*SuperPang* works by first homogenizing the input sequences so that homologous regions come to have identical sequences, and then build a pangenome assembly graph. In this graph, homologous regions are dereplicated and linked together following their synteny in the input genomes, providing a transparent and useful representation of the pangenome architecture. The regions of the pangenome that were always found together in the input genomes will be represented as non-branching paths (NBPs), with branching points indicating either reordering events or interruption of synteny due to the presence of accessory genes. NBPs thus naturally identify modules of genes that preserve synteny and are always inherited or transferred together, potentially due to functional relatedness (e.g. gene operons or genomic islands).

The mOTUpan algorithm (Buck *et al*., 2021) is then run to classify each NBP into the core and accessory genomes based on their occurrence in the input sequences. The NBPs are then exported as nucleotide sequences, which facilitates a modular analysis of the core and accessory genomes of the species. Furthermore, the graph is traversed in order to combine the different NBPs into longer contigs that follow the consensus gene ordering in the pangenome.

The result of running *SuperPang* is a non-redundant reference assembly for the input species which, unlike the representative genomes obtained by dereplication methods such as drep (Olm *et al*., 2017), contains both the core and accessory genomes. This assembly can be annotated with tools such as Prodigal (Hyatt *et al*., 2010), Prokka (Parks, 2014) or SqueezeMeta (Tamames &Puente-Sánchez, 2019) in order to identify protein-coding sequences, RNA-genes or other non-coding elements of interest, but it will also preserve the information on the intergenic regions, potentially allowing for the discovery of new non-coding, regulatory regions involved in intra-species diversification. This assembly will unambiguously recruit virtually all the shotgun meta’omic reads originating from the input species, enabling variant calling not only in the core, but also within the accessory genome. Read mapping can also be easily used to track the prevalence of the different accessory genes across populations. The *SuperPang* assembler is thus a versatile tool for high-resolution studies of intra-species diversity. Below, we provide details on its implementation and performance, as well as several examples highlighting its potential applications.

## SOFTWARE IMPLEMENTATION AND AVAILABILITY

*SuperPang* is implemented in the Python3 and Cython3 programming languages and released under the GNU General Public License v3.0. The source code is available at https://github.com/fpusan/SuperPang. *SuperPang* can be easily installed using *pip* (https://pypi.org/project/SuperPang/) or *conda* (https://anaconda.org/fpusan/superpang).

## BENCHMARKING

Execution time, resource utilization, and overall performance were evaluated by running *SuperPang* on 113 previously published genomes from *Polynucleobacter paneuropeus* (Hoetzinger *et* al., 2021; **Supplementary Table S1**), a pelagic proteobacterial species abundant in humic lakes and ponds, with genome sizes around 1.8 Mbp. The availability of multiple sequenced isolates from a broad geographic range, sharing >96.5% ANI and an open pangenome make it a suitable model for benchmarking the software. We ran SuperPang using 24 cores on an increasingly large number of genomes, and we collected statistics on the total execution time, maximum memory usage, the total size in base pairs of the predicted core and accessory genomes, and the completeness and contamination of the predicted core genome (**Figure 2; Supplementary Table S2**).

**Figure 2.**
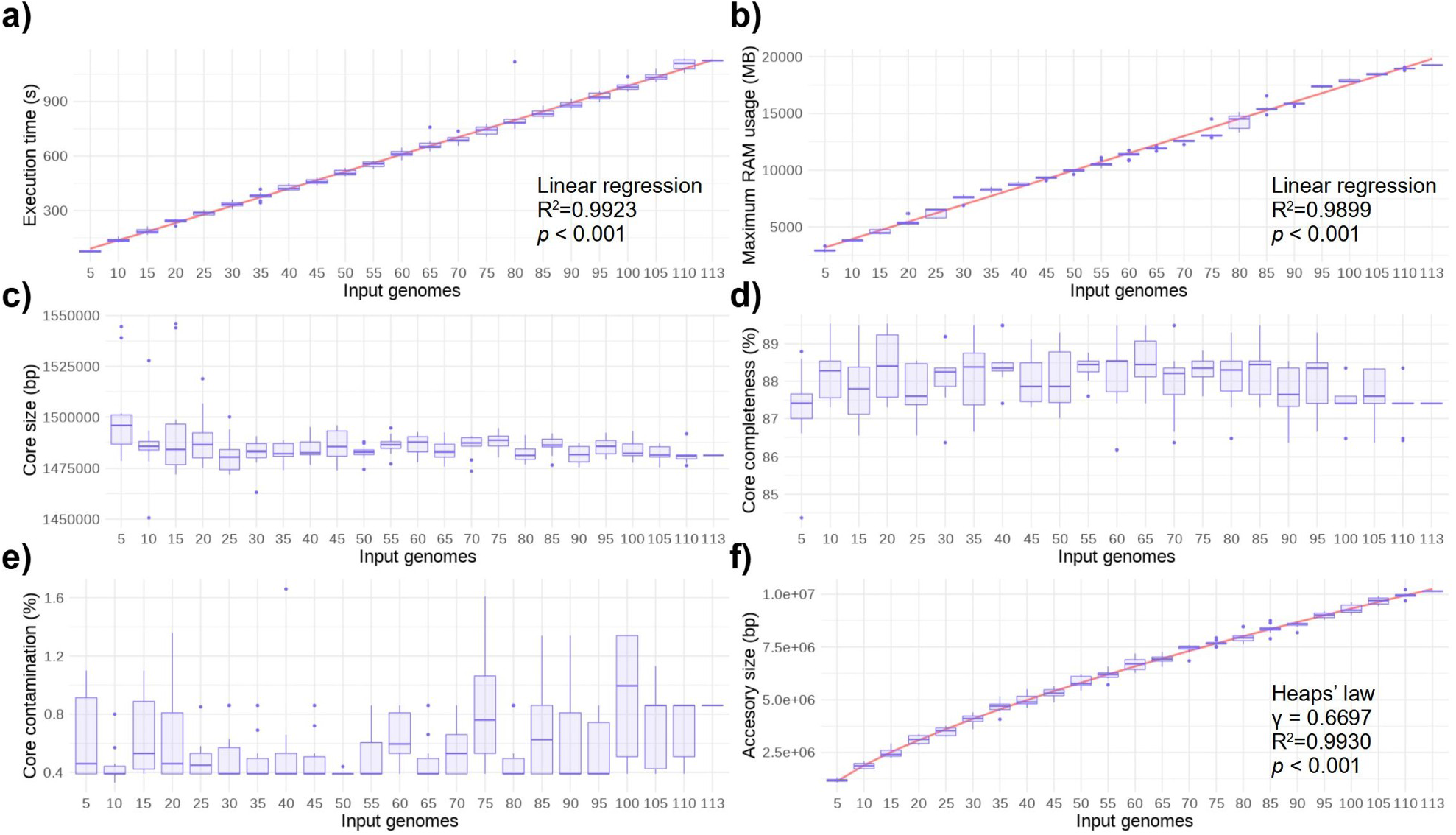
Benchmarking of *SuperPang* with 113 input genomes. For an increasing number of input genomes (10 replicates) the boxplots show the evolution of **a)** execution time, **b)** maximum memory usage, **c)** estimated core genome size, **d)** estimated core genome completeness, **e)** estimated core genome contamination, **f)** estimated accessory genome size.

Execution time increased linearly with the number of input genomes enabling rational upscaling to larger datasets (**Figure 2a**; linear model, adjusted R^2^ = 0.9899, p < 0.001). Memory usage also increased linearly with the number of input genomes, up to 19.9 Gb for 113 genomes (**Figure 2b;** linear model, adjusted R^2^ = 0.9929, p < 0.001). The size of the predicted core genome had a median of 1.49 Mbp for 5 input genomes, after which it decreased slightly and stabilized around 1.48 Mbp (**Figure 2c**). Core completeness remained stable around 88% with the number of input genomes (**Figure 2d**), while core contamination, which acts as a proxy for the core duplication level, remained around 1% (**Figure 2e)**. This verifies earlier findings that the mOTUpan algorithm yields an accurate prediction of the core genome, even when the number of input genomes is low (Buck *et al*., 2021). In contrast, the size of the predicted accessory genome kept increasing with the number of input genomes (**Figure 2f**). Heap’s law regression (Tettelin *et al*., 2008) on the accessory size versus the number of input genomes gave a γ parameter of 0.67, suggesting that *Polynucleobacter paneuropeus* has an open pangenome (**Figure 2f**, power law regression, adjusted R^2^ = 0.993, p < 0.001), as previously concluded (Hoetzinger *et* al., 2021).

We further tested the completeness and redundancy of the *SuperPang* assembly by generating synthetic short reads from the 113 input genomes and mapping them to the *assembly*.*fasta* file produced by *SuperPang*. The mapping percentage was of 96%, while only 5% of the reads mapped to more than one position. Our results show that *SuperPang* was able to successfully collapse the homologous regions present in the 113 input genomes and predict a complete and non-redundant core genome for the species while still capturing the totality of the accessory genome in the assembly.

### TEST CASE 1: IDENTIFICATION OF GENOMIC ISLANDS

We first reconstructed the pangenome of *Polynucleobacter asymbioticus*, which analogous to *P. paneuropeus* is a widespread planktonic freshwater bacterium with a dynamic flexible genome hosting a diverse set of mobile genomic islands (Hoetzinger *et al*., 2017). Running *SuperPang* on genomes from nine isolates of the species yielded a core genome of 1.75 Mbp and 216 accessory NBPs longer than 1000 bp (1.44 Mbp in total). The majority of them (214, 1.44 Mbp) assembled into the main scaffold together with the core NBPs, which makes them good candidates for representing genomic islands. We then assessed if the accessory part of the *SuperPang* assembly could recover genomic islands (GIs) that were defined in an earlier and more laborious approach as consecutive sequences of auxiliary genes longer than 10 kbp. Indeed, all 28 GI variants identified in the previous study were recovered, and >50% of the accessory genome of each strain was covered by NBPs longer than 10 kbp (compare **Figure 3a** with Figure 3 in Hoetzinger *et al*., 2017). Aligning core and accessory NBPs to reference genomes provide a quick method to reveal GIs present in individual genomes and open for further studies to gain insights about their function and origin (**Figure 3b**). This alignment illustrates two recombination hot spots within the *P. asymbioticus* genomes, characterized by large gaps within the core genome where multiple accessory NBPs align to. These regions were described earlier to represent replacement GIs, which are typically found at conserved genomic locationsand often contain hypervariable sequences that were suggested to be transferred between genomes through recombination that utilizes sequence homology at their bounderies (López-Pérez *et al*., 2013; López-Pérez *et al*., 2014). NBPs that only align through small stretches of their sequences to certain reference genomes might indicate fragments transferred this way, where the aligned stretches might represent homologous sequences potentially utilized for recombination. Examples of such NBPs in **Figure 3b** are 5, 34 and 3 aligning to QLW-P1DMWA-1 or 26 and 17 to MWH-RechtKol4, respectively. The recombination hot spot (**Figure 3b**) that has been referred to as CSC (cell surface composition) previously (Hoetzinger *et al*., 2017) is a prime example of a replacement GI. Annotation suggests that many genes within this region are involved in cell surface glycosylation. Similar genome regions seem to be ubiquitous in prokaryotic genomes and the gene repertoire present in such genomic islands was proposed to determine the strain glycotype, i.e. a decisive feature for phage recognition (López-Pérez *et al*., 2016). NBPs obtained from the *SuperPang* assembly that are present in the respective genome regions (NBP 6 for QLW-P1DMWA-1 and NBPs 1 and 8 for MWH-RechtKol4, see **Figure 3b**) could provide means to trace such glycotypes in metagenomes.

Another type are so called additive GIs, which are associated with mobile genetic elements and can harbor a variable number of gene cassettes that are typically flanked by integrases or transposases and/or tRNAs that may serve as target sites for integrases (López-Pérez *et al*., 2013). Six gene cassettes located within additive GIs were associated to metabolic functions in the previous study of *P. asymbioticus* (Hoetzinger *et al*., 2017), namely assimilatory nitrate reduction (NIT), lipid metabolism (LIP), carbon monoxide dehydrogenation (COD), sulfate transport (SUL) and heavy metal resistance (HMR). While we recovered these gene cassettes from the accessory NBPs, some of them were split into multiple NBPs due to sequence divergence among the reference genomes (**Figure 3c**). This drawback can be overcome by using the consensus contigs present in the *assembly*.*fasta* file, which combine NBPs according to their synteny in the sequence graph. By splitting this assembly at the core-accessory boundaries, accessory contigs were obtained that contained complete functional gene cassettes (**Figure 3d, Supplementary Figure S2**). These results confirm that partitioning the auxiliary genome into contigous sequences rather than separated genes, as done by conventional tools for pangenome analysis (Page *et* al., 2015; Gautreau *et al*., 2020), carries biological meaning in line with the emergence principle.

**Figure 3.**
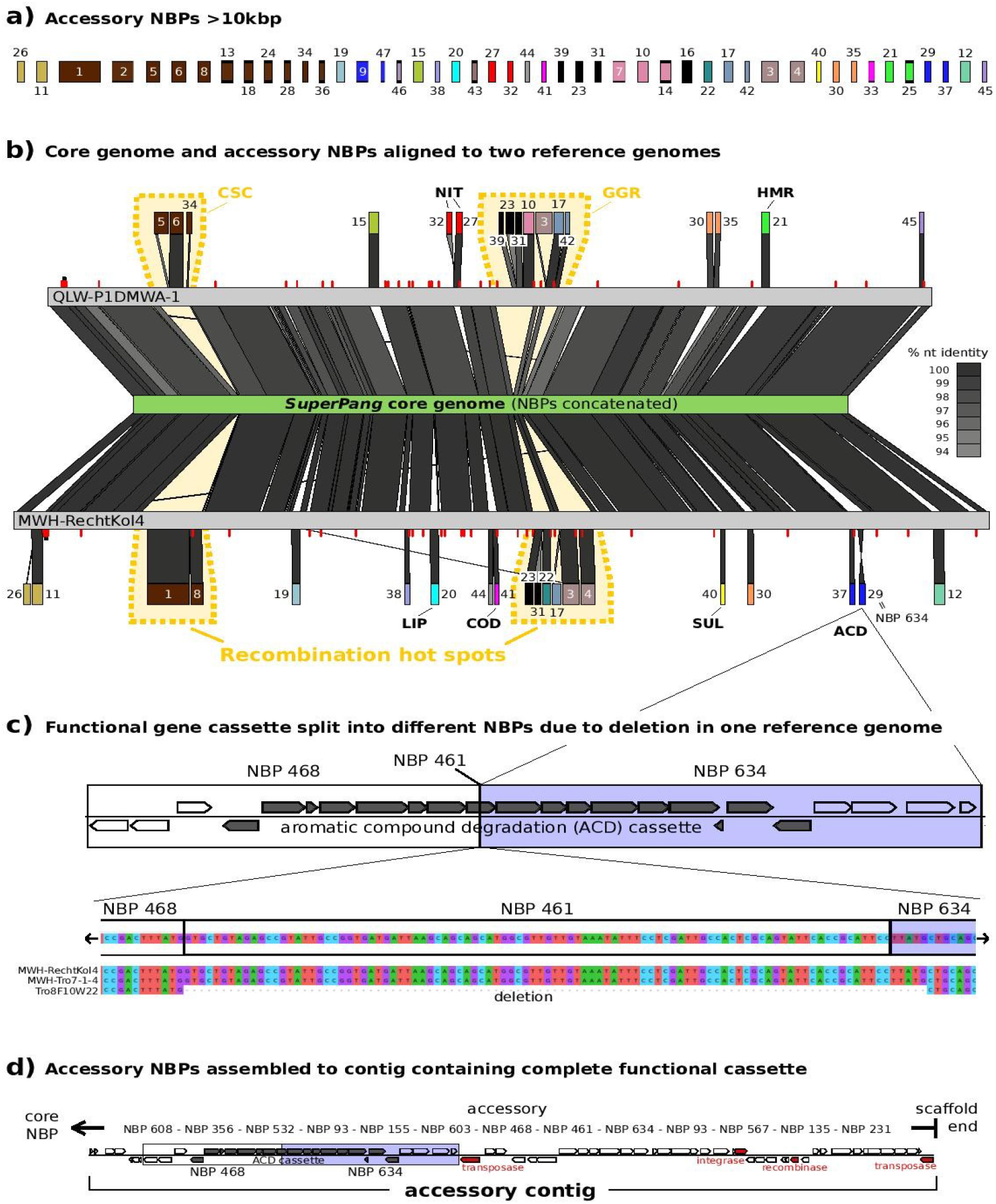
Illustrating genomic islands in reference genomes using the SuperPang assembly. **a)** Accessory NBPs longer than 10 kbp obtained from *SuperPang*. The NBPs are colored according to Figure 3 in Hoetzinger *et al*. (2017) after alignment against the *P. asymbioticus* genomes. They are numbered in descending order of sequence length and arranged according to the order they appear in the reference genomes. **b)** Core NBPs were ordered against the genome of strain QLW-P1DMWA-1 and concatenated. The thereby obtained core genome is represented in the middle, and blastn hits to two reference genomes are depicted in grayscale according to sequence identity. GIs appear as alignment gaps, i.e. sequences present in a reference genome that do not align to the core. Red bars indicate tRNAs of the reference genomes. Blastn alignments to the auxiliary NBPs as shown in a) are displayed above and below QLW-P1DMWA-1 and MWH-RechtKol4, respectively. NBPs aligned to GIs for which a specific function has been inferred previously Hoetzinger *et al*. (2017) are named (CSC, cell surface composition; NIT, assimilatory nitrate reduction; HMR, heavy metal resistance; LIP, lipid metabolism; COD, carbon monoxide dehydrogenation; GGR, giant gene region; SUL, sulfate transport; ACD, aromatic compound degradation). **c)** Example of a functional gene cassette split into separate NBPs, caused by a 101 bp long deletion in a reference genome. Genes associated to the ACD cassette are highlighted by grey fill color. The nucleotide sequence of the NBPs in the split region is shown together with an alignment of the respective sequence in three reference genomes. **d)** Accessory contig obtained from the SuperPang assembly containing the complete ACD cassette. NBPs contained in the contig are given above the gene representation.

### TEST CASE 2: EXTENDING PANGENOMES WITH ENVIRONMENTAL SEQUENCES IN KNOWN MICROBIAL SPECIES

We sought to generate an extended pangenome assembly of *P. paneuropaeus* that included novel diversity from environmental sequences. For this, we screened StratFreshDB (Buck *et al*., 2021), a dataset shotgun reads and assemblies from more than 250 environmental freshwater samples, and retrieved the MAGs that shared more than 95% average nucleotide identity (ANI) to any of our *Polynucleobacter asymbioticus* genomes. This search recovered 5 MAGs with less than 5% contamination and variable completeness levels (from 87% to 9%; **Supplementary Table S1**). We combined these MAGs with the isolate genomes into a new dataset, and ran *SuperPang* to generate a new assembly that, using the isolate genomes as a backbone, could also incorporate new information that was only present in the environmental MAGs. This allowed us to identify 40 new accessory NBPs (296 kb in total) longer than 1000 bps that assembled into the main scaffold. Among them, we found what appeared to be a prophage that was missing in the isolate genomes, but present in the MAGs (**Figure 4; Supplementary Table S3**). The graphical nature of the *SuperPang* output allowed us to easily determine the genomic context where the phage had integrated (**Figure 4c**). Finally, 5 accessory NBPs (13 kb in total) were found only in the two low-quality MAGs (9.97 % and 9.34 % completeness respectively), but nonetheless assembled into the main scaffold. This shows the potential of *SuperPang* to recover information also from incomplete assemblies coming from environmental metagenomes, provided that other more complete genomes are also available.

**Figure 4.**
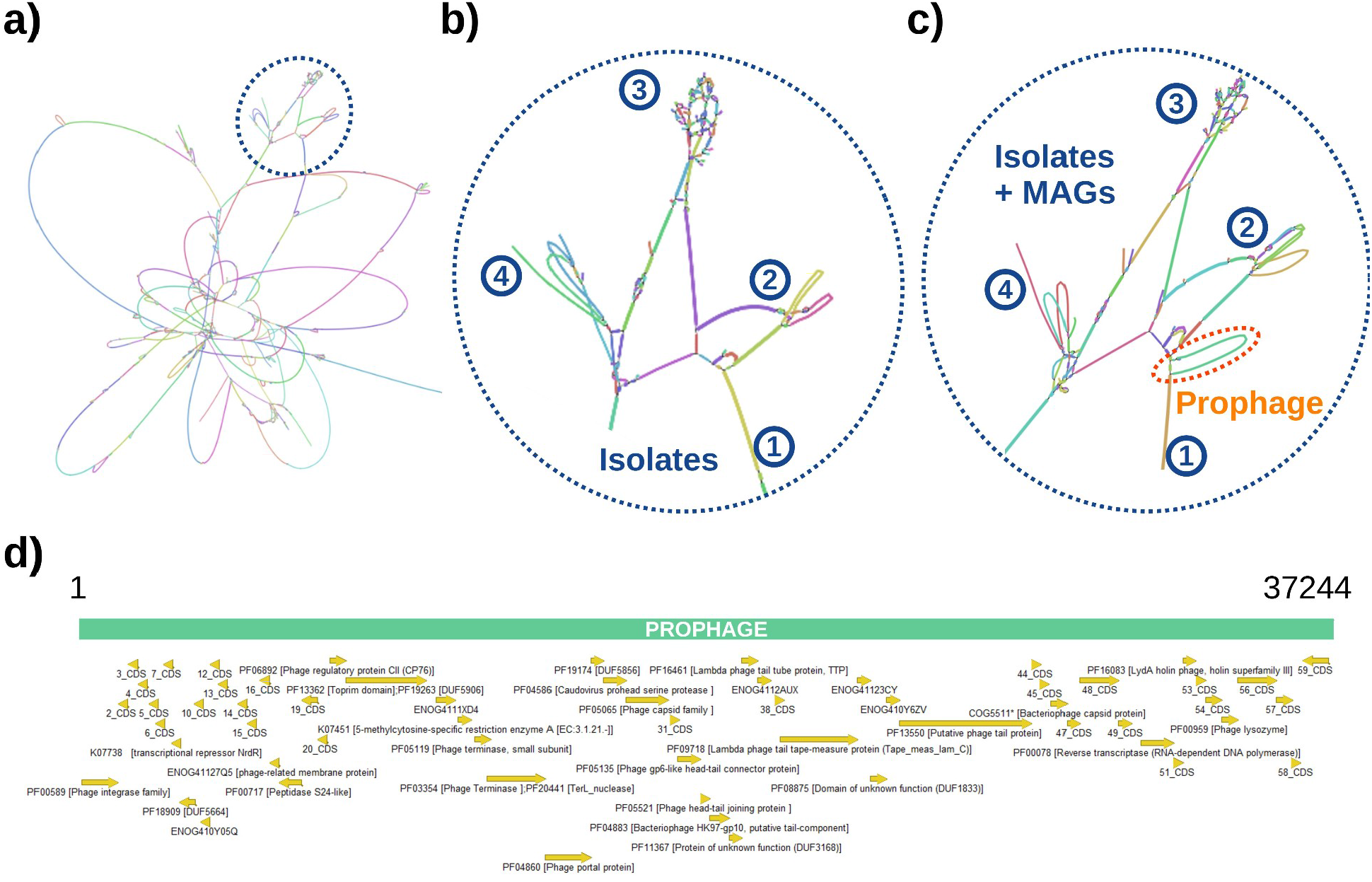
Extending pangenomes with environmental sequences. **a)** Pangenome assembly obtained from 9 *Polynucleobacter asymbioticus* isolates. Insets show the same area before (center) and after (right) adding the information from 5 additional metagenome-assembled genomes (MAGs). Numbers indicate homologous regions in both assemblies. A non-branching path appearing only in the MAGs and corresponding to a prophage is marked in orange. **b)** genetic map of the environmental prophage sequence.

### TEST CASE 3: SPATIOTEMPORAL DYNAMICS OF THE CORE AND ACCESSORY GENOMES IN UNCULTIVATED MICROORGANISMS

We ran *SuperPang* on 44 metagenome-assembled genomes (MAGs) from the same *Polynucleobacter* species, and used the results to analyze its population structure over a set of 44 samples (7 time points, variable depths) from lake Loclat (Switzerland). We considered two sources of intra-species diversity: point mutations leading to allelic variation and the gain/loss of accessory genes, which leads to differences in gene content between populations. This allowed us to study the intra-species dynamics of the core and accessory genomes in both time and space.

We first studied population structure by performing an ordination of the samples based on their pairwise fixation index (F_ST_), which measures population differentiation in terms of allelic frequencies such that more dissimilar populations will have a higher pairwise F_ST_ value. We calculated F_ST_ indices for the core and the accessory genomes separately, and visualized them via Principal Coordinates Analysis (PCoA). For the core, samples clustered by date, with a sharper transition between the first time point and all the others (**Figure 5a**). This transition corresponds in time with the fall mixing of the lake, which occurs after the first time point (**Supplementary Figure S1**). A similar pattern could be observed for the accessory genome, although it was less clear (**Figure 5b**).

**Figure 5.**
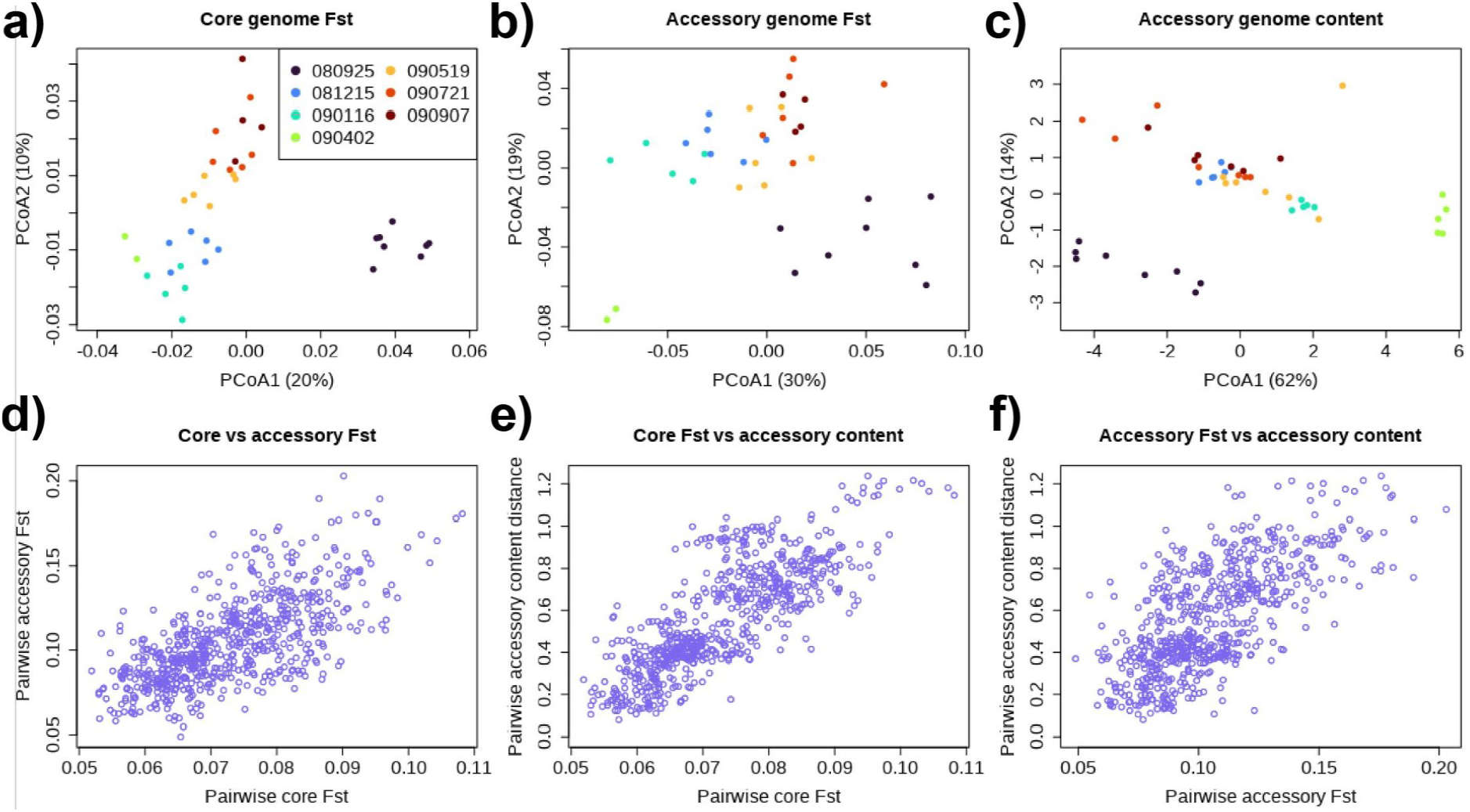
Characterizing the core and accessory genome dynamics of uncultivated species. **Upper row:** Principal Coordinate Analysis (PCoA) ordinations showing the distribution of samples from lake Loclat (Switzerland) according to the population structure of an uncultivated *Polynucleobacter* species. Samples are colored based on the sampling date. Ordinations were performed based in **a)** core genome FST (i.e. population differentiation based on allele frequencies), **b)** accessory genome FST and **c)** accessory gene content (i.e. population differentiation based on the prevalence of accessory genes. **Lower row:** comparison between different metrics for assessing population differentiation. Each point represents a pair of samples. **d)** comparison of core genome FST and accessory genome FST. **e)** comparison of core genome FST and distance in accessory genome content. **f)** comparison of accessory genome FST and distance in accessory genome content.

We then tracked the dynamics of the accessory NBPs across our samples to assess how accessory gene abundances change under different selective pressures. As our input genomes were uncurated MAGs, there was a risk that some of the input sequences were contaminants coming from other species, even after we had selected only the MAGs with less than 3% contamination according to CheckM. To further mitigate this risk, we only focused on the accessory NBPs that assembled into the main scaffold and were thus connected to the core genome in the assembly graph. Finally, we discarded the NBPs that could not be assigned to the *Polynucleobacter* genus, or had too low or too high average copy numbers per genome. This left us with a set of 65 high-confidence NBPs totaling 244997 bp (33% of the unfiltered accessory genome size). From these data, we generated a matrix of the dissimilarity between pairs of *Polynucleobacter* populations based on their accessory gene content, and visualized it using PCoA (**Figure 5c**).

Samples from the first time point clustered separately from the rest (**Figure 5c**, black points), as previously observed for the core and accessory F_ST_. Samples from December, January and April formed very tight clusters (**Figure 5c**, blue, turquoise and green points), as opposed to the rest of the time points, where samples were more scattered. Interestingly, the former coincided with the period when the water column of lake Loclat was subject to wind-driven homogenization (**Supplementary Figure S1**). The existence of such tighter clusters suggest that, when the lake is mixed, the species is strongly dominated by a single population. Similarly, the higher dissimilarities in accessory gene content when the lake is stratified may imply the existence of different subpopulations being successful at different depths.

We also assessed whether differences in allelic frequencies (F_ST_) could be predictive of the accessory gene content. Our results suggest that this is the case, showing a high agreement between core F_ST_, accessory F_ST_ and accessory gene content dissimilarity (**Figure 5d,e,f**). This correlation was however noisy, suggesting that accessory genes might not be always bound to certain alleles in the core genome. Finally, we investigated how the prevalence of individual NBPs in the population changed with time or in response to environmental variables (**Figure 6**). Some of the NBPs presented a sharp transition in copy number after the first sample (**Figure 6a,b,c**), similar to the one observed for allelic frequencies in the core genome (**Figure 5a**). We could however also find examples of NBPs whose prevalence was associated with temperature, dissolved sulfate, or dissolved nitrate (**Figure 6d,e,f**), highlighting the flexibility and complexity of environmental factors influencing prokaryotic pangenomes. While further discussion is beyond the scope of this manuscript, our results show how pangenome assemblies produced by *SuperPang* can be used to track the core and auxiliary genome dynamics of wild populations of bacteria, even if they come from uncultivated species.

**Figure 6.**
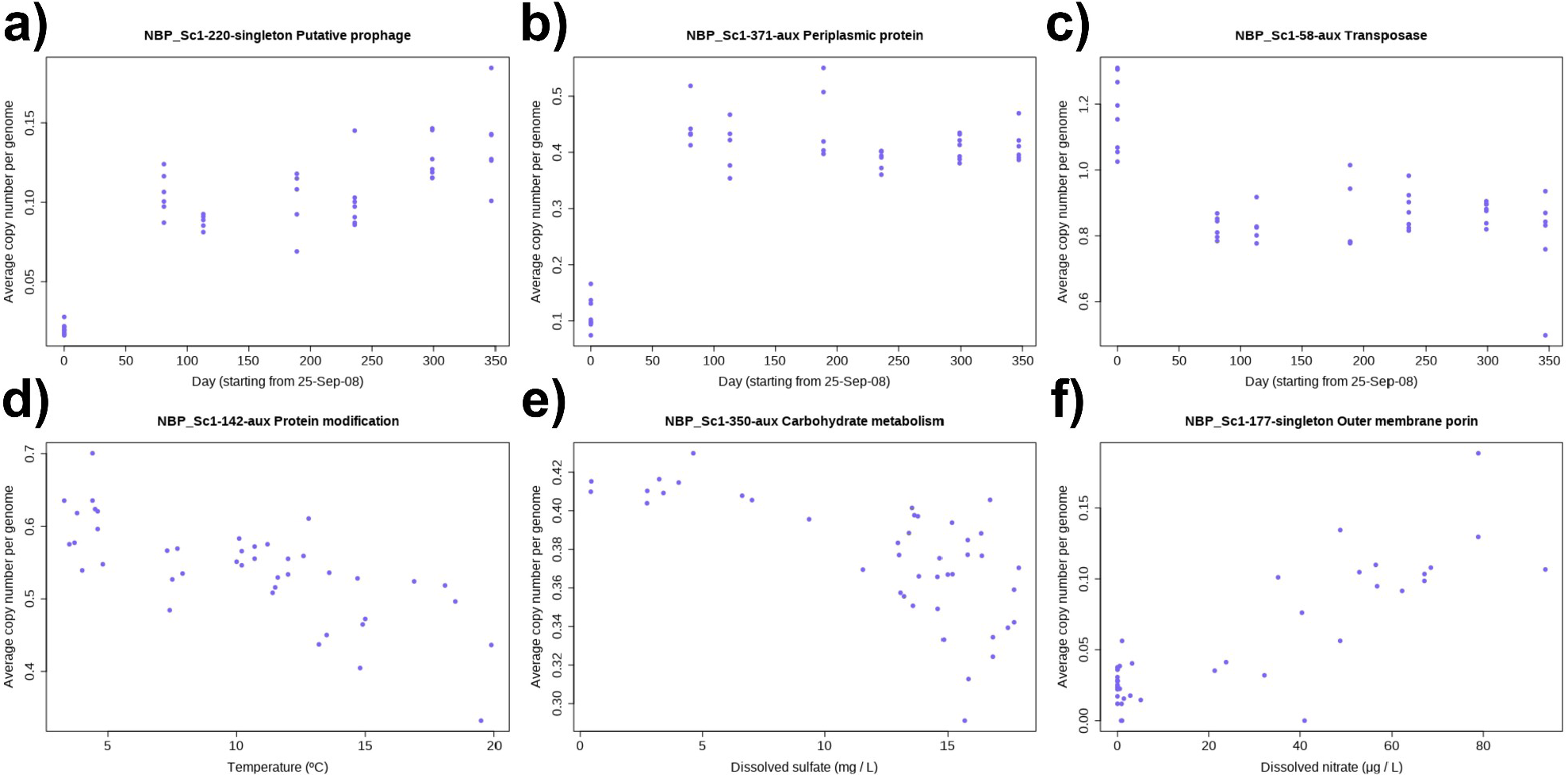
Tracking the response of individual genomic islands to environmental change. Examples of accessory regions in the pangenome of an uncultivated *Polynucleobacter sp*. Inhabiting lake Loclat (Switzerland), whose prevalence in the population correlates with time or with different environmental variables. All examples shown have an adjusted R^2^ > 0.3 and a FDR < 0.01 according to a linear model. The predicted proteins and functions for each NBP can be found in **Supplementary Table S4**.

## CONCLUSIONS

*SuperPang* produces pangenome assemblies from a set of input genomes of variable quality, including metagenome-assembled genomes. The results from *SuperPang* are complete (contain the combined core and accessory genomes), non-redundant (each orthologous region appears only once), preserve gene ordering and contain both coding and non-coding regions. Notably, S*uperPang* does not attempt to resolve individual strains but rather provides a modular view of the pangenome by reporting the regions that are always inherited together (here called non-branching paths, or NBPs). Our approach naturally identifies operons and genomic islands, and enable tracking of their prevalence in different populations of the species. This will enable high-resolution studies of microbial diversity and facilitate simultaneous analysis of allelic and gene content variation between multiple populations of the same species and as such take us one step further to describe and study the finer details of naturally complex microbial communities.

## ACKNOWLEDGEMENTS

Computational resources were provided by the Department of Aquatic Sciences and Assessment (Swedish University of Agricultural Sciences) and by resources in projects SNIC 2021/5-53 and SNIC 2021/22-602 provided by the Swedish National Infrastructure for Computing (SNIC) at UPPMAX, partially funded by the Swedish Research Council through grant agreement no. 2018-05973. FPS was supported by the European Union’s Horizon 2020 research and innovation programme under the Marie Skłodowska-Curie grant agreement No 892961. MH was supported by Formas project 2019-02336 as part of the ERA-NET BlueBio Cofund project ImprovAFish. SB and MB were supported by the Swedish Research Council project 2017-04422.

## METHODS

### Computational resources

All analyses were performed in an Ubuntu 20 workstation with an AMD Ryzen 9 5900X 12-Core Processor and 128 Gb of RAM. When applicable, processes were ran using 24 CPU threads. Data used for benchmarking were stored in a 1Tb Samsung SSD 980 PRO 1TB drive. Preliminary testing was also performed in UPPMAX (Uppsala Multidisciplinary Center for Advanced Computational Science) computing cluster.

### Input data

*SuperPang* accepts a set of FASTA files, each containing the contigs from a different genome. Genomes are assumed to belong to the same phylogenetic cluster (e.g. same species, or same 95% ANI mOTU). Genomes can be complete genomes, draft genomes, MAGs or SAGs. Optionally, the user can also supply the standard output obtained by running CheckM (Parks *et al*., 2015) over the input genomes. The completeness of each input genome will then be used to differentiate between the core and accessory contigs in the final assembly (see below).

### Homogenization of input sequences

Prior to assembly, we identify homologous regions in the different input sequences and homogenize them so that they will later assemble into the same contig. The objective is to generate a pangenome assembly that contains the entirety of the core and accessory genomes, but in which polymorphisms are collapsed into a single consensus sequence, thus avoiding duplications in the assembled core genome. Homogenization is performed by the following iterative algorithm:

1 While the number of iterations is less than a certain threshold *r* and at least one sequence was homogenized sequence in the previous iteration.
  1.1 Sort the input sequences by decreasing length.
  1.2 For each *target* sequence, visited in decreasing length order, that has not been corrected by a longer target sequence during this iteration.
    1.2.1 For each *query* sequence smaller than the *target* sequence, that has not been corrected by a longer target sequence during this iteration, and has not been corrected by this target sequence in previous iterations.
      1.2.1.1 Align *target* and *query* using *minimap2* (Li, 2018) with the following parameters *-Hk19, -w5, -e0, -m100, --rmq=yes, --dual=no, -DP, --no-long-join, - U50,500, -g10k, -s200*.
        1.2.1.1.1 Let *M* be the number of matches and *X* be the number of mismatches. If *M / (M+X)* exceeds a certain threshold *i*, homogenize *query* with *target*. For each non-match CIGAR operation in the *query-target* alignment relating a segment in the *query* target to a segment in the *target* sequence, if the length of the operation is below a certain threshold (*m* for matches, *g* for indels), replace that contiguous segment in the *query* sequence by its counterpart in the *target* sequence.
        1.2.1.1.2 Mark *query* as a homogenized by *target* in this iteration.
  1.3 Write the sequences (including any changes resulting from homogenization) to a temporary output file to be used as the input for the next iteration.
2 Use the output of the last iteration as the final output.

The different thresholds can be controlled by the user, with defaults of *i*=0.95, *m*=100, *g*=100, *r*=20. Increasing *m* and *g* will result in a more aggressive homogenization: longer contiguous mismatch and indel stretches will be homogenized, reducing the duplication levels in the core genome after pangenome assembly, but potentially also removing some accessory genes. Sequence homogenization is performed by default when running *SuperPang*, but can also be run independently by calling the *homogenize*.*py* standalone script.

### Pangenome assembly

Input sequences are stored into a De-Bruijn graphs (DBG) using Graph-Tool (Peixoto, 2014). Non-branching paths (NBP) in the DBG are then used to build a sequence graph (SG), in which each node represents a NBP path and vertices join NBPs that have been observed to overlap in the same input sequence. The SG is then split into connected components, and pairs are identified such that the sequences from both members are each-other’s reverse complements. For each pair, the component with the highest numbers of NBPs in the same orientation as the input sequences is deemed the forward component and reported as a scaffold in the assembly. NBP coverage is defined as the average prevalence of its constituent kmers in the input genomes, we note that this is unrelated to the coverage of the NBP in any particular sample. NBPs sharing the first and last kmers (i.e. corresponding to a bubble in the DBG) will be aligned pairwise using the *mappy* module from the *minimap2* suite (Li, 2018). In case the homology between them is higher than a certain threshold *b* (default 0.95) only the longest NBP will be kept in the assembly.

### Core genome identification

NBPs are classified with the bayesian classifier mOTUpan (Buck *et al*., 2021) in order to determine their likelihood to belong to the core or the accessory genome. Sequences sharing a fraction *a* of its kmers (default *a*=0.5) with an input genome are deemed to be present in that genome. For each NBP sequence, mOTUpan then classifies it as core/accessory/singleton by taking into account its prevalence in the input genomes, and the estimated completeness of those genomes (Buck *et al*., 2021). If completeness estimates are not provided as an input, SuperPang assumes a 50% starting completeness and lets mOTUpan calculate posterior estimates.

### Contig generation

NBPs are combined by traversing the SG with the following greedy algorithm.

1. Calculate the origins of the SG (i.e. the NBPs with no predecessors).
2. While there are valid origins,

1. Extend each origin (sorted by decreasing coverage) by a deep first search, selecting the successors with the highest coverage. The same NBP can be visited twice within the same extension, but not if it was visited when extending a different origin.
2. Remove the visited nodes from the SG and recalculate its origins.

The assembler outputs a FASTA file containing the contigs, and a FASTG file (following the SPAdes / MEGAHIT specification) containing assembly graph and suitable for visualization in Bandage (Wick *et al*., 2015). A separate info file keeps track of which regions of each contig were deemed to be core, accessory or singletons. Individual NBPs are also outputted into three different files containing all the NBPs, the core NBPs, and the accessory NBPs respectively.

### Benchmarking

Execution time, resource utilization, and overall performance were evaluated by running *SuperPang* (v0.9.2) on 113 previously published *Polynucleobacter paneuropeus* genomes (**Supplementary Table S1**). *SuperPang* was run using 24 CPU threads on an increasing number of genomes, in multiples of five, until finally running it in the totality of the 113 genomes. For each step, *SuperPang* was run in ten replicates by drawing the requested number of genomes at random without replacement. For each *SuperPang* run, we tracked the total execution time, maximum memory usage, the total size in base pairs of the predicted core and accessory genomes, and the completeness and contamination of the predicted core genome as measured by CheckM (Parks *et al*., 2015) using the marker genes for the genus *Polynucleobacter*).

1 million synthetic metagenomic reads of 125 bp were simulated from the unassembled 113 input genomes using InSilicoSeq (Gourlé *et al*., 2019). The *--mode perfect and --coverage uniform* parameters were used to generate reads with no errors that covered the input genomes uniformly. Reads were mapped back to the *SuperPang* assembly obtained from the same 113 input genomes using blastn (Altschul, 1990) with a threshold of 95% identity and 90% query coverage. We then counted the proportion of reads that mapped at least once to the SuperPang assembly, and also the proportion of reads that mapped in two or more positions.

Pangenome openness was evaluated by fitting a power law of the form S ∼ N^γ^, where S is the predicted accessory genome size and N is the number of input genomes, where the exponent γ > 0 indicated an open pangenome (Tettelin *et al*., 2008).

### Recovery of Metagenome Assembled Genomes (MAGs) from Polynucleobacter asymbioticus

StratFreshDB (Buck et al., 2021) was screened for MAGs that were taxonomically assigned to *P. asymbioticus*, which yielded seven MAGs with >10% completeness and <3% contamination according to checkm (Parks *et al*., 2015). ANI values between these MAGs and the genomes of *P. asymbioticus* isolates were computed and only MAGs that shared >95% ANI with all genomes of isolates were kept, resulting in five MAGs ranging from 12 – 78% completeness and 0 – 0.9% contamination.

### Annotation of genomic islands

Genomic islands were functionally annotated with SqueezeMeta/SQMtools (Tamames &Puente-Sánchez, 2019; Puente-Sánchez *et al*., 2020). Briefly, ORFs were identified with Prodigal (Hyatt *et al*. 2010) and annotated against the KEGG (Kanehisa &Goto, 2000) and EggNog4.5 (Huerta-Cepas *et al*., 2016) databases using DIAMOND (Buchfink *et al*., 2015). Additionally, hmmer3 (Eddy, 2009) was used to annotate ORFs against the PFAM database (Finn *et al*., 2010). APE (https://jorgensen.biology.utah.edu/wayned/ape/) was used for visualization.

### Core and accessory dynamics of environmental species

We ran *SuperPang* (v0.9.2) on genomes from a *Polynucleobacter* metagenomic Operational Taxonomic Unit (mOTU) that was previously published as part of the StratFreshDB (Buck *et al*., 2021). In total, 44 metagenome-assembled genomes (MAGs) with completeness above 40% and contamination below 3% were selected (**Supplementary Table S1**). We then tracked the dynamics of the core and accessory NBPs (contained in the *NBPs*.*core*.*fasta* file generated by *SuperPang*) over a set of 44 metagenomes corresponding to a time-series study of the lake Loclat (Switzerland; seven time points, different depths; see Buck *et al*., 2021).

We first used POGENOM 0.8.3 (Sjöqvist *et al*., 2021) to calculate population genomics parameters for the species in relation to the set of samples. For the *Input_POGENOM* pre-processing pipeline, parameters were kept as default, except for *mode_prefilt* which was set to FALSE. The *pogenom*.*pl* script was run with *–subsample TRUE* and --*min_count* 10. The --*min_found* parameter was set to 1 for calculating FST indices, and to 37 for calculating nucleotide diversity and non-synonymous to synonymous mutation ratios. FST indices were reported as the average of 100 *pogenom*.*pl* runs, and used directly for Principal Coordinate Analysis (PCoA).

We then used SqueezeMeta (Tamames and Puente-Sánchez, 2019) to taxonomically and functionally annotate the NBPs present in the *NBPs*.*fasta* file generated by *SuperPang*. The raw abundances of the NBPs and their average copy-number per genome in the different samples were estimated as described in Puente-Sánchez et al. (2020), using a set of 10 single-copy marker genes previously described in Salazar *et al*., 2019.

For calculating the differences in gene content between populations and minimize the effect of possible contaminant contigs in our input MAGs, we selected the NBPs that 1) were classified as accessory, 2) had more than 1000 bases, 3) had an average copy-number per genome between 0.1 and 1.5, 4) had at least one ORF assigned to the *Polynucleobacter* genus by SqueezeMeta and 5) assembled into the main scaffold. The NBPs in the main scaffold are either the core genome or accessory regions directly connected to the core genome in the assembly graph. Focusing on the main scaffold thus provides extra safety against the presence of contaminant contigs in MAGs. Differences in gene content between populations were visualized using PCoA on a matrix of euclidean distances that was obtained after centered log-ratio (CLR) transformation of the raw NBP abundances.

## SUPPLEMENTARY MATERIAL

**Supplementary Figure S1**. Oxygen concentration across time and depth in a time series from lake Loclat (Switzerland).

**Supplementary Figure S2**. Analogous to Figure 3a/3b but using accessory contigs obtained from the *SuperPang* assembly instead of NBPs.

**Supplementary Table S1**. Genomes used in this work

**Supplementary Table S2**. Benchmarking of the SuperPang assembler on 113 *Polynucleobacter paneuropeus* genomes

**Supplementary Table S3**. Annotation of a putative prophage non-branching path in the

*Polynucleobacter asymbioticus* pangenome

**Supplementary Table S4**. Annotation of six non-branching paths in a Polynucleobacter sp. pangenome

**Supplementary Data S1**. Genome sequence of *Polynucleobacter asymbioticus* T8_bin-239.

**Supplementary Data S2**. Genome sequence of *Polynucleobacter asymbioticus* T8-022_bin-141.

