## Supplementary figures and images for "Exploring intra-species diversity through non-redundant pangenome assemblies"

### Supplementary Figure S1

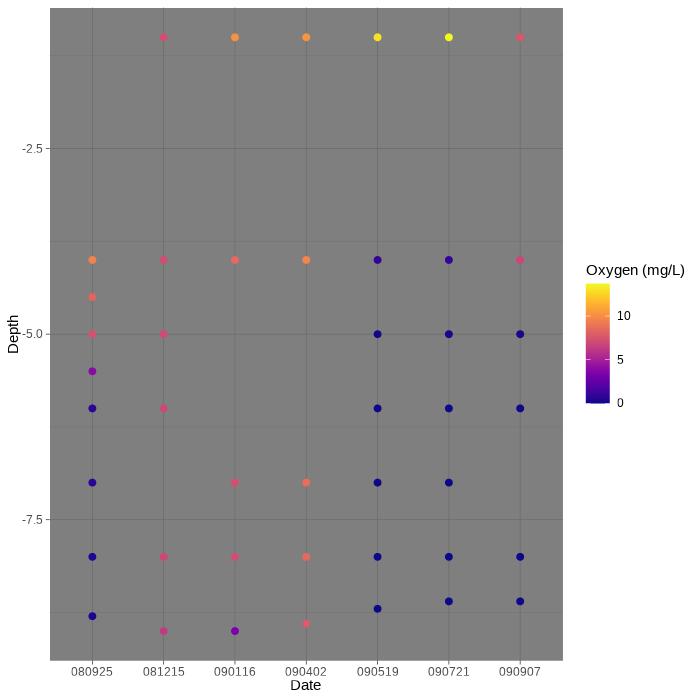

### Supplementary Figure S2

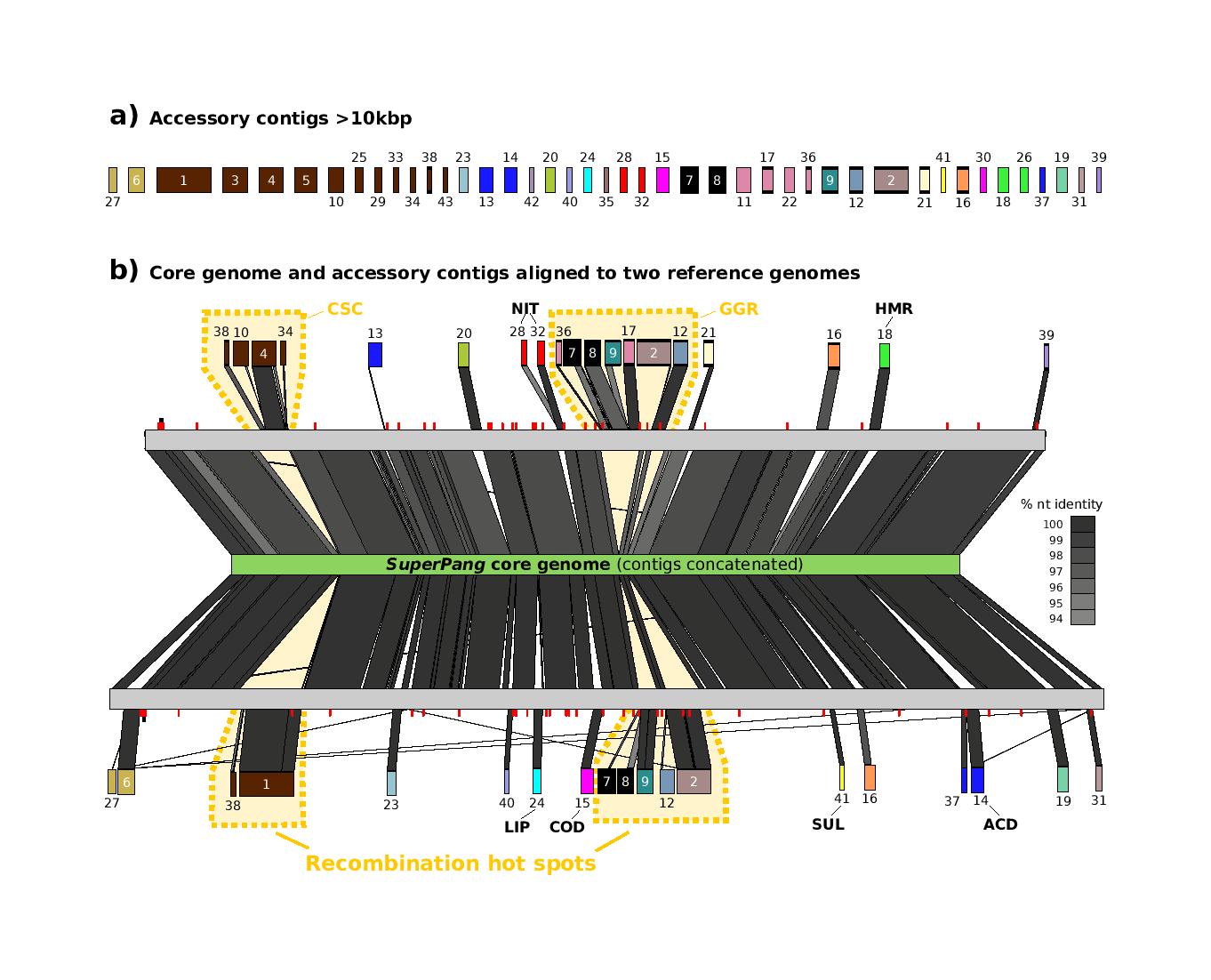
